# An mRNA expression-based signature for oncogene-induced replication-stress

**DOI:** 10.1101/2021.02.19.431930

**Authors:** Sergi Guerrero Llobet, Arkajyoti Bhattacharya, Marieke Everts, Bert van der Vegt, Rudolf S.N. Fehrmann, Marcel A.T.M. van Vugt

**Author notes:** shared first author (Sergi Guerrero Llobet, Arkajyoti Bhattacharya). shared senior authorship (Rudolf S.N. Fehrmann, Marcel A.T.M. van Vugt). correspondence; Marcel van Vugt, PhD.

## Abstract

**Background:** Oncogene-induced replication stress characterizes many aggressive cancers, including triple-negative breast cancer (TNBC). Several drugs are being developed that target replication stress, although it is unclear how tumors with high levels of replication stress can be identified. We aimed to develop a gene expression signature of oncogene-induced replication stress.

**Methods:** TNBC and non-transformed RPE1-*TP53*^wt^ and RPE1-*TP53*^mut^ cell lines were engineered to overexpress the oncogenes *CDC25A*, *CCNE1* or *MYC*. DNA fiber analysis was used to measure replication kinetics. Analysis of RNA sequencing data of cell lines and patient-derived tumor samples (TCGA n=10,592) was used to identify differential gene expression. Immunohistochemical validation was conducted on breast cancer samples (n=330).

**Results:** RNA sequencing revealed 52 commonly upregulated genes after induction of *CDC25A*, *CCNE1* or *MYC* in our cell line panel. Integration with gene expression data of TGCA samples with amplification of replication stress-inducing oncogenes (*CDC25A*, *CCNE1*, *MYC*, *CCND1*, *MYB*, *MOS*, *KRAS*, *ERBB2*, and *E2F1*), yielded a six-gene signature of oncogene-induced replication stress (*NAT10*, *DDX27*, *ZNF48*, *C8ORF33*, *MOCS3,* and *MPP6)*. Expression of NAT10 in breast cancer samples was correlated with phospho-RPA (R=0.451, p=1.82×10^−20^) and γH2AX (R=0.304, p=2.95×10^−9^). Applying the oncogene-induced replication stress signature to patient samples (TCGA n=8,862 and GEO n=13,912) defined the replication stress landscape across 27 tumor subtypes, and identified diffuse large B cell lymphoma, ovarian cancer, TNBC and colorectal carcinoma as cancer subtypes with high levels of oncogene-induced replication stress.

**Conclusion:** We developed a gene expression signature of oncogene-induced replication stress, which may facilitate patient selection for agents that target replication stress.

## INTRODUCTION

A subset of cancers, including triple-negative breast cancer (TNBC) shares a profound genomic instability^1,2^ This phenomenon is characterized by continuous gains and losses of chromosomal fragments and complex genomic rearrangements. As a consequence, genomic instability underlies the rapid acquisition of genomic aberrations that drive therapy failure^3^. Finding new treatment options for genomically unstable cancers is not only relevant for TNBC, but also other hard-to-treat cancers with extensive genomic instability, including high-grade serous ovarian cancer (HGSOC)^4^. Genomic instability is observed in multiple aggressive cancer subtypes and is associated with the inability of cancer cells to faithfully repair DNA damage^5^. Increasingly, an important source of the DNA lesions that fuels genomic instability appears to be replication stress^6,7^.

During S-phase of the cell cycle, all DNA must be replicated in a coordinated manner, which is initiated at genomic loci called ‘replication origins’^8^. Replication origins are fired in a temporally-controlled way, which prevents exhaustion of the nucleotide pool and warrants the availability of essential components of the replication machinery^9^. DNA replication can be challenged in various ways, which is collectively referred to as replication stress. An important cause of replication stress is the uncoordinated initiation of origin firing due to oncogene activation^7,10,11^. Consequently, oncogene activation leads to depletion of the nucleotide pool and collisions between of the replisome and the transcription machinery, resulting in slowing or complete stalling of replication forks^7,9,12^. When stalled replication forks are not resolved in time, they can collapse and cause DNA double-strand breaks (DSBs) and lead to genomic instability.

Several oncogenes have been linked to induction of replication stress, many of them leading to aberrant activation of CDK2. For instance, overexpression of the CDK2-binding partner Cyclin E1 (*CCNE1*) or the CDK2-activating phosphatase CDC25A leads to replication slow-down and reversal of replication forks^13^. In line with this notion, amplification of *CCNE1* has been proposed as a biomarker for tumors with high levels of replication stress^14^. However, multiple other oncogenic events beyond *CCNE1* amplification also can lead to replication stress, including overexpression of *MYC*^15,16^, *MOS*^17^, *E2F1*^*11*^, or expression of the E6/E7 HPV oncoproteins^9^. Currently, there is no uniform way to determine oncogene-induced replication stress levels in cancer samples.

Identification of cancers with high levels of replication stress is increasingly relevant because several drugs have been developed that specifically target tumor cells with high levels of replication stress. Inhibition of WEE1 has been shown to have therapeutic efficacy in HGSOC, which is considered as a prototypical tumor type with high levels of replication stress^18^. Interestingly, patients with the largest decrease in tumor size upon WEE1 inhibitor treatment showed enrichment for *CCNE1* amplification^18^. In line with this notion, overexpression of *CCNE1* sensitized TNBC cell lines to WEE1 inhibition^14,19^. Furthermore, an unbiased genomic screen identified regulators of CDK2 as determinants of WEE1 inhibitor sensitivity^20^. Mechanistically, WEE1 inhibition is thought to exacerbate levels of replication stress further, while it inactivates the G2/M cell cycle checkpoint, driving cells into mitotic catastrophe^21^. Similarly, inhibition of the ATR or CHK1 checkpoint kinases has been shown to preferentially target tumor cells with molecular characteristics that are associated with replication stress^22–24^.

It is currently unclear how patients can be optimally selected for treatment with agents that target replication stress. To this end, we performed gene expression profiling of cell lines with oncogene-induced replication. Further refinement with expression profiles of patient derived tumor samples yielded a gene expression signature of replication stress, which allowed us to describe a pan-cancer landscape of oncogene-induced replication stress.

## RESULTS

### Overexpression of CDC25A, CCNE1, or MYC results in replication stress

To develop an mRNA expression-based signature for oncogene-induced replication stress, we engineered a panel of cell lines to overexpress CDC25A, CCNE1, or MYC in a doxycycline-inducible manner. This cell line panel included non-transformed human retina epithelial (RPE1) cell lines, either with wild type *TP53* (RPE1-*TP53*^wt^) or a derivative in which *TP53* was mutated using CRISPR-Cas9 (RPE1-*TP53*^mut^), and TNBC cell lines MDA-MB-231, BT549 and HCC-1806 (Fig. 1A, Suppl. Fig. 1). To validate that oncogene induction indeed affected DNA replication, DNA fiber analysis was performed (Fig. 1B). A severe reduction in DNA synthesis velocity was observed upon induction of CDC25A, CCNE1, or MYC, as measured by IdU fiber tract lengths in RPE1-*TP53*^wt^ cells (32%, 23% and 42% decrease in CDC25A, CCNE1, and MYC overexpressing cells versus controls, respectively, Fig. 1C). Similar decreases in DNA replication dynamics were observed in RPE1-*TP53*^mut^ cells (34%, 27%, and 51% decrease in CDC25A, CCNE1 and MYC overexpressing cells versus controls, respectively, Fig. 1D), indicating that these effects are independent of *TP53* status. Subsequently, the impact of oncogene expression on DNA synthesis was analyzed in TNBC cell lines. Again, we consistently observed shortening of IdU tract lengths in MDA-MB-231, BT549 and HCC-1806 cell lines upon doxycycline-induced overexpression of CDC25A, CCNE1 or MYC (Fig. 1E-G), but not in empty vector controls (Fig. 1C-G). Taken together, these data indicate that induction of CDC25A, CCNE1, or MYC results in replication stress in cancer cells, as well as in untransformed cell lines, independently of *TP53* status.

**Figure 1:**
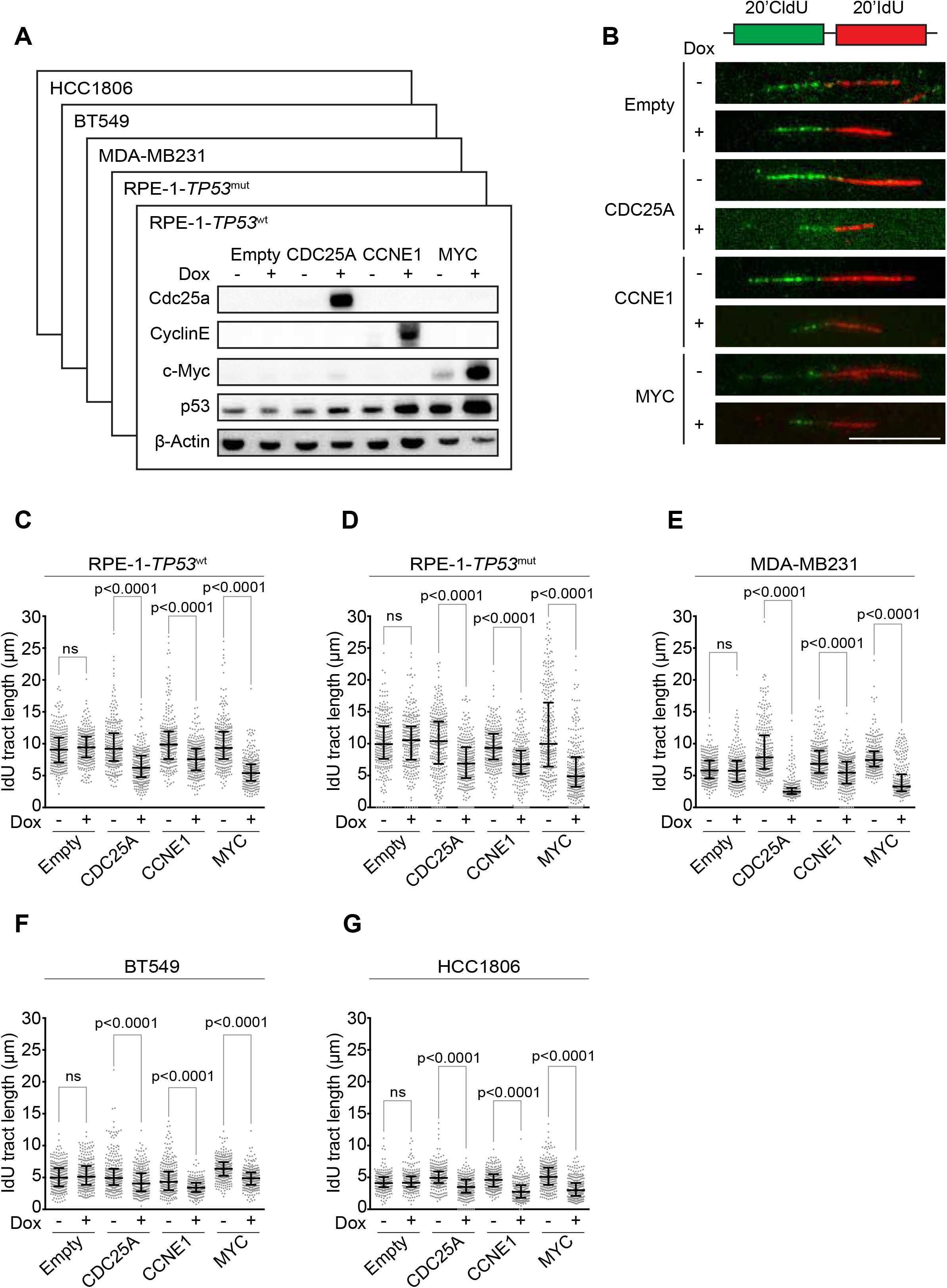
Overexpression of CDC25A, CCNE1 or MYC leads to replication stress. **(A)** Indicated RPE1-*TP53*^wt^ cell lines were treated with doxycycline for 48 hours. Immunoblotting was performed for CDC25A, CCNE1, p53, and β-Actin. **(B)** Cells were treated with doxycycline as described for panel A and subsequently pulse-labeled for 20 minutes with CldU (25 μM) and 20 minutes with IdU (250 μM). Representative DNA fibers from RPE1-*TP53*^wt^ cells are shown. Scale bar represents 10 μm. **(C-G)** Quantification of IdU DNA fiber lengths, as described in panel B. Per condition, 300 fibers from RPE1-*TP53*^wt^ cells were analyzed. P values were calculated using the Mann-Whitney U test for RPE1-*TP53*^wt^ (panel C), RPE1-*TP53*^mut^ (panel D), MDA-MB-231 (panel E), BT549 (panel F) and HCC1806 (panel G).

### Gene expression profiling of cell line models with oncogene-induced replications stress

To map the transcriptional consequences of overexpression of CDC25A, CCNE1 and MYC, RNA sequencing of RPE1-*TP53*^wt^, RPE1-*TP53*^mut^, MDA-MB-231, BT549, and HCC-1806, cells was performed both at 48 hours and at 120 hours after oncogene induction to capture gene expression changes provoked by replication stress (Fig. 2A). Subsequently, all cell lines and genetic perturbations were analyzed in a pooled fashion. To this end, the pooled RNAseq dataset was first normalized to remove any possible effects of doxycycline treatment in control cell lines as well as cell line-specific effects (see Supplementary Methods for details). Subsequently, permutational multivariate analysis of variance (PERMANOVA) was performed to identify genes that were differentially expressed upon overexpression of each oncogene (CDC25A, CCNE1, and MYC) across cell lines (Fig. 2B). Induction of CCNE1 or CDC25A led to significantly differentially upregulated genes (n=1,330 and n=309, respectively; p<0.01), with only 2 genes being downregulated (Fig. 2A, right panel). In contrast, MYC overexpression resulted in a substantial differential gene expression, involving both downregulation (n=2,576) and upregulation (n=935) of gene expression (Fig. 2A, right panel). Interestingly, expression of 52 genes was found to be commonly upregulated in response to induction of CCNE1, CDC25A, or MYC (Fig. 2A, right panel).

**Figure 2:**
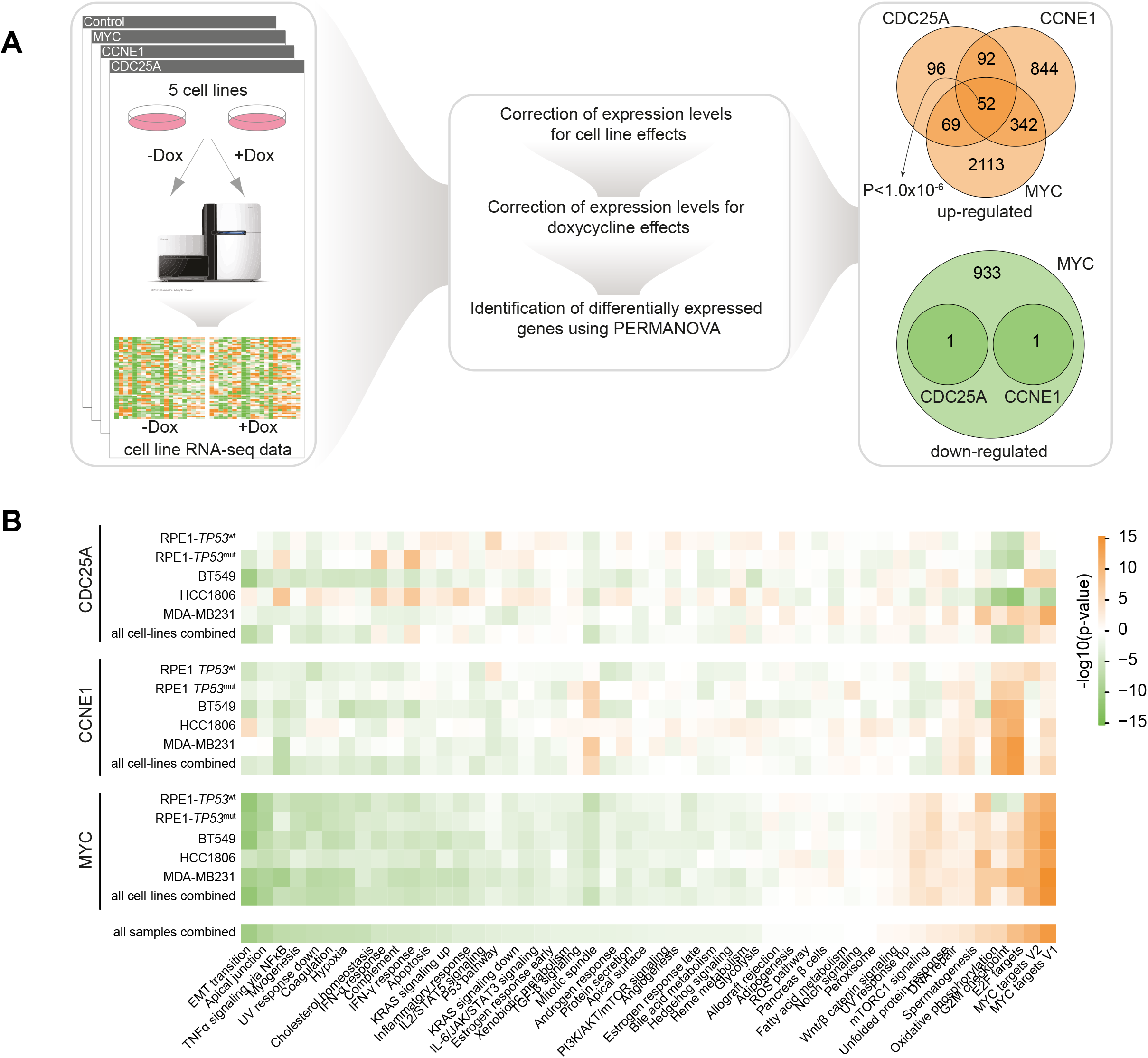
Overexpression of CDC25A, CCNE1, or MYC leads to upregulation of 52 genes. **(A)** RNAseq data of 5 cell lines were corrected for cell line-specific and doxycycline treatment effects on gene expression. Subsequently, PERMANOVA was used to identify common differentially expressed genes upon oncogene expression. 52 genes were found to be commonly upregulated whereas no genes were found to be commonly downregulated in response to induction of CCNE1, CDC25A or MYC **(B)** Gene Set Enrichment Analysis using two-sided Welch’s t-test for the MSigDB Hallmark collection. An orange box indicates enrichment for upregulated genes due to overexpression of oncogenes in corresponding cell lines, and a green box indicates enrichment for downregulated genes.

Importantly, gene-set enrichment analysis (GSEA) on metrics obtained from PERMANOVA (see Supplementary Methods for details) revealed that common biological pathways were affected upon induction of CCNE1, CDC25A, or MYC (Fig. 2B), with strong upregulation in expression of MYC targets, and genes involved in cell cycle control and oxidative phosphorylation (Fig. 2B). In contrast, expression of genes related to biological pathways involved in cellular morphology and inflammatory signaling were commonly downregulated (Fig. 2B). Taken together, these results indicate that oncogene overexpression induces distinct yet overlapping gene expression changes, affecting common biological pathways.

### Common differential gene expression upon oncogene overexpression between in vitro models and patient samples

To investigate whether the differential gene expression we observed in our cell line models overlaps with patient derived tumor samples with amplification of these oncogenes, we retrieved copy number data and mRNA expression data from The Cancer Genome Atlas (TCGA) (Fig. 3A). TCGA samples were classified into two groups based on the copy number of each of the oncogenes (’amplified’ versus ‘neutral’). After that, genes that were significantly differentially expressed upon amplification of each oncogene (CDC25A, CCNE1, or MYC) were identified (Fig. 3A). Upon amplification of the three oncogenes, expression of 720 common genes was significantly upregulated (permutation test: p<1.0×10^−6^), and that of 597 genes was down-regulated (permutation test: p<1.0×10^−6^). GSEA revealed strong upregulation in expression of genes related to MYC targets, cell cycle control, and oxidative phosphorylation (Fig. 3B). In contrast, expression of genes related to immune signatures was commonly downregulated in samples with oncogene amplification (Fig. 3B). The majority of the enrichments in the TCGA data were similar to those obtained by differential expression analysis from the cell line data. Of note, two genesets (i.e., “allograft rejection” and “IL-6/JAK/STAT3 signaling”) were only significantly enriched in the analysis with patient derived tumor samples. This is in line with immune activities in the patient samples not being reflected in cell line models.

**Figure 3:**
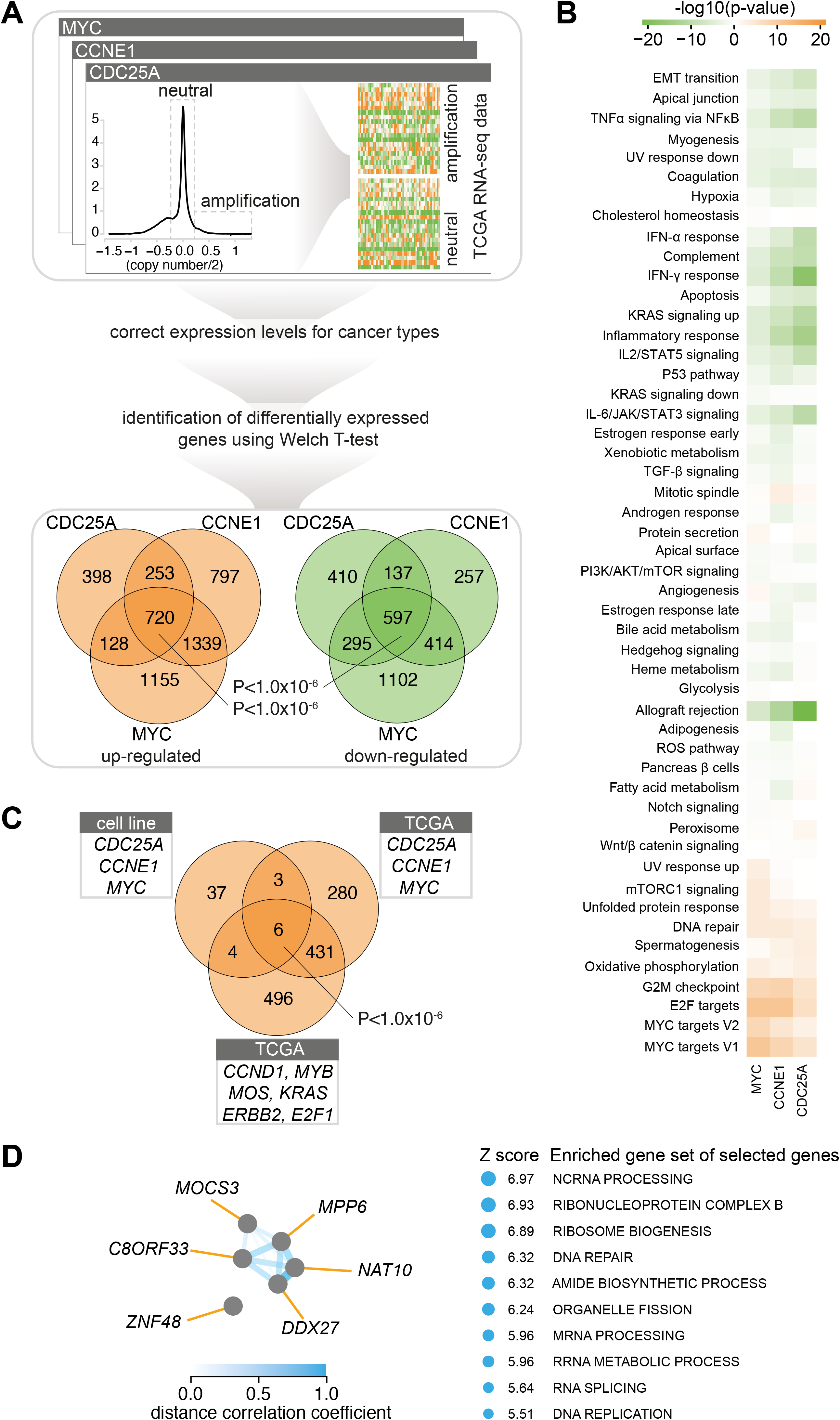
Common differential gene expression of 6 genes upon oncogene overexpression between *in vitro* models and patient samples. **(A)** TCGA RNAseq samples were queried for amplification of *CCNE1*, *CDC25A* or *MYC*. Expression of 720 genes was found to be commonly upregulated, whereas 597 genes were found to be commonly downregulated in response to amplification of CCNE1, CDC25A or MYC. **(B)** Gene Set Enrichment Analysis using two-sided Welch’s t-test for the MSigDB Hallmark collection. An orange box indicates enrichment for upregulated genes due to overexpression of oncogenes in corresponding cell lines, whereas a green box indicates enrichment for downregulated genes. **(C)** TCGA RNAseq samples were queried for amplification of *CCND1*, *MYB*, *MOS*, *KRAS*, *ERBB2* and *E2F1.* The resulting commonly upregulated genes were overlaid with upregulated genes identified upon overexpression of CCNE1, CDC25A or MYC in cell lines and upregulated genes in TCGA samples with *CCNE1*, *CDC25A* or *MYC* amplifications, resulting in 6 commonly upregulated genes **(D)** Co-functionality analysis of commonly upregulated genes using the GenetICA algorithm.

Since replication stress can also be caused by oncogenes beyond *MYC*, *CCNE1*, and *CDC25A*, we additionally identified commonly upregulated genes in TCGA tumor samples overexpressing other oncogenes that have been associated with replication stress, including *CCND1*, *MYB*, *MOS*, *KRAS*, *ERBB2,* and *E2F1*^11,17,25–30^. (Fig. 3C). Analysis of genes that were commonly upregulated upon oncogenes expression in cell line models as well as patient-derived tumor samples revealed six genes (i.e., *NAT10*, *DDX27*, *ZNF48*, *C8ORF33*, *MOCS3*, and MPP6) (Fig. 3C, D). Co-functionality analysis of commonly upregulated genes using the GenetICA algorithm (see Supplementary Methods for details) pointed at roles for these genes in ncRNA processing, DNA repair, and ribosome biogenesis (Fig. 3D).

### NAT10 protein expression is associated with markers of replication stress in breast cancer samples

To validate the mRNA-based replication stress signature, we selected NAT10 for immunohistochemical analysis. The acetyltransferase NAT10 (N-Acetyltransferase-10) has previously been implicated in regulation of various processes, including regulation of the DNA damage response^31,32^ and regulation of translation^33,34^. NAT10 is a nuclear protein, with predominant localization to the nucleolus^35^. Immunohistochemical analysis of NAT10 staining in a series of breast cancer tissues (n=410), confirmed nuclear localization of NAT10, with nucleolar expression in a subset of cancer samples (Fig. 4A). Importantly, a significant variation in protein expression was observed, with triple-negative breast cancer samples showing the highest NAT10 expression levels (Fig. 4B). Of note, absence or presence of nucleolar localization was not different between breast cancer subgroups (Fig. 4B).

**Figure 4:**
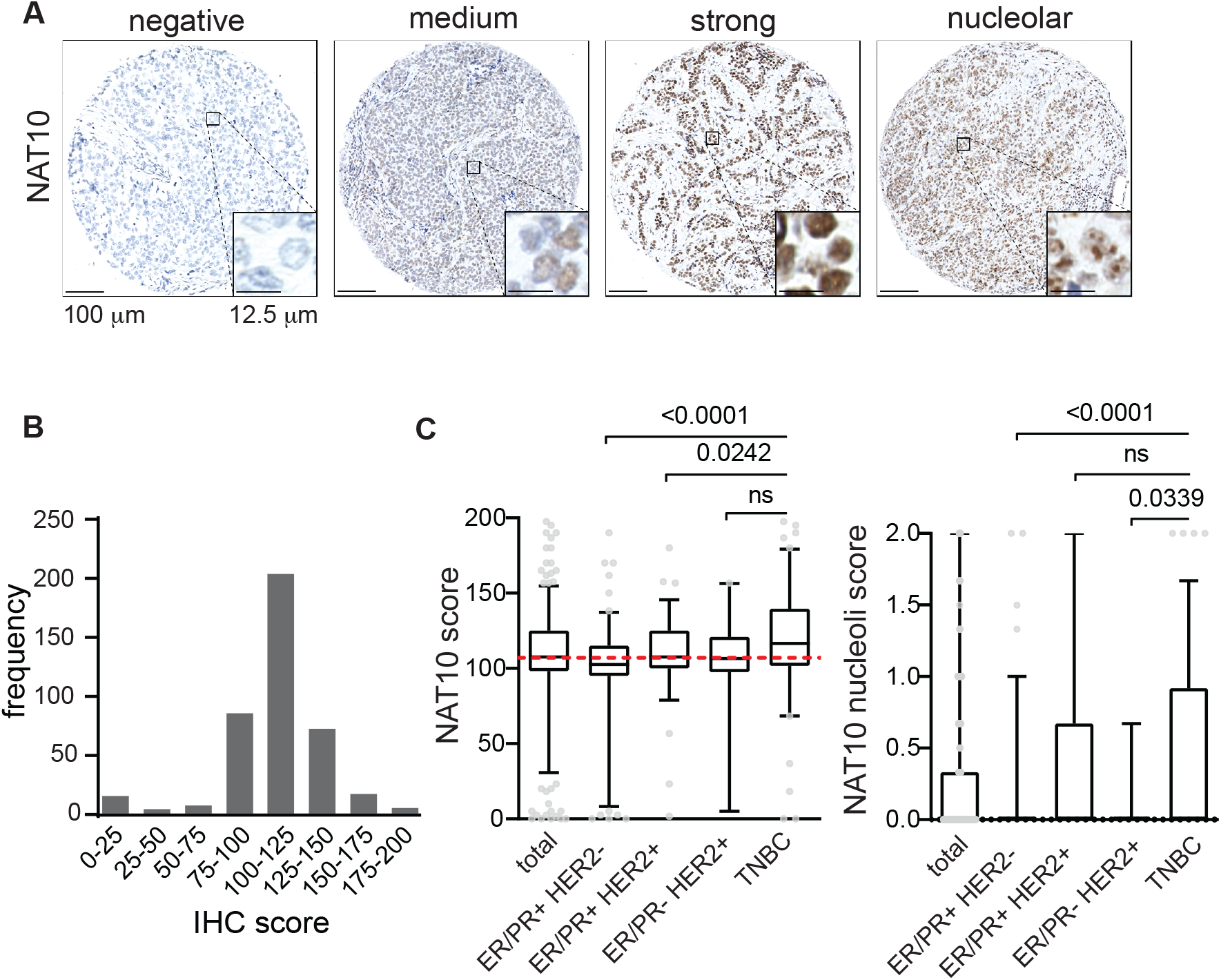
Immunohistochemical analysis of NAT10 in breast cancer patients. **(A)** Representative staining of NAT10 in breast cancer patient samples (n=410). Scale bar represents 100 μm. **(B)** Patients from the combined cohort (n=410) and breast cancer subgroups ER/PR^+^HER2^−^ (n=164), ER/PR^+^HER2^+^ (n=95), ER/PR^−^HER2^+^ (n=21) and TNBC (n=130) were analyzed. Tumor tissue was immunohistochemically scored for expression of NAT10 and NAT10 nucleoli **(C)**, indicated P values were calculated using Mann-Whitney U test. Interquartile ranges are displayed, dashed red line represents the median score from the combined cohort and outliers are shown as dots.

**Figure 5:**
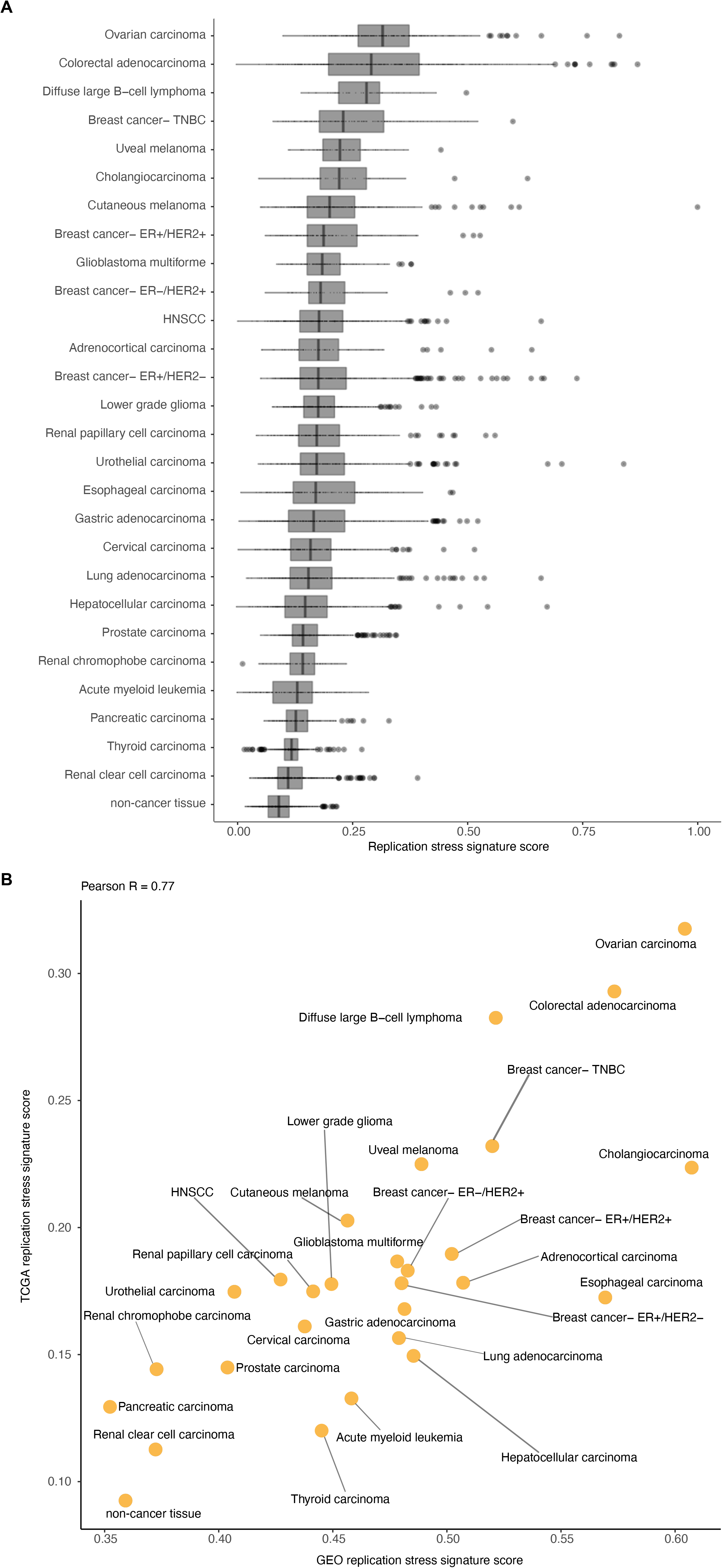
The landscape of oncogene-induced replication stress across cancer types. **(A)** Distribution of the replication stress signature across 27 cancer types as well as non-cancerous tissues of patient samples from TCGA. **(B)** Scatter plot of replication stress signature for 27 cancer types as well as non-cancerous tissues of patient samples from TCGA (y axis) and GEO (x axis).

For this breast cancer cohort, we previously reported the presence of p-RPA, a marker of replication stress, and γH2AX, which reflects DNA breaks, a possible result of stalled replication forks collapse^36^. In line with NAT10 expression being part of the oncogene-induced replication stress signature, Spearman correlation analysis showed an association of NAT10 expression with both p-RPA (R=0.451, p=1.82×10^−20^) and γH2AX (R=0.304, p=2.95×10^−9^) (Table 1A). Subgroup analysis showed that the most significant correlations between NAT10 expression and pRPA or γH2AX were observed in ER/PR^−^/HER2^+^ and TNBC samples (Table 1A).

**Table 1:**
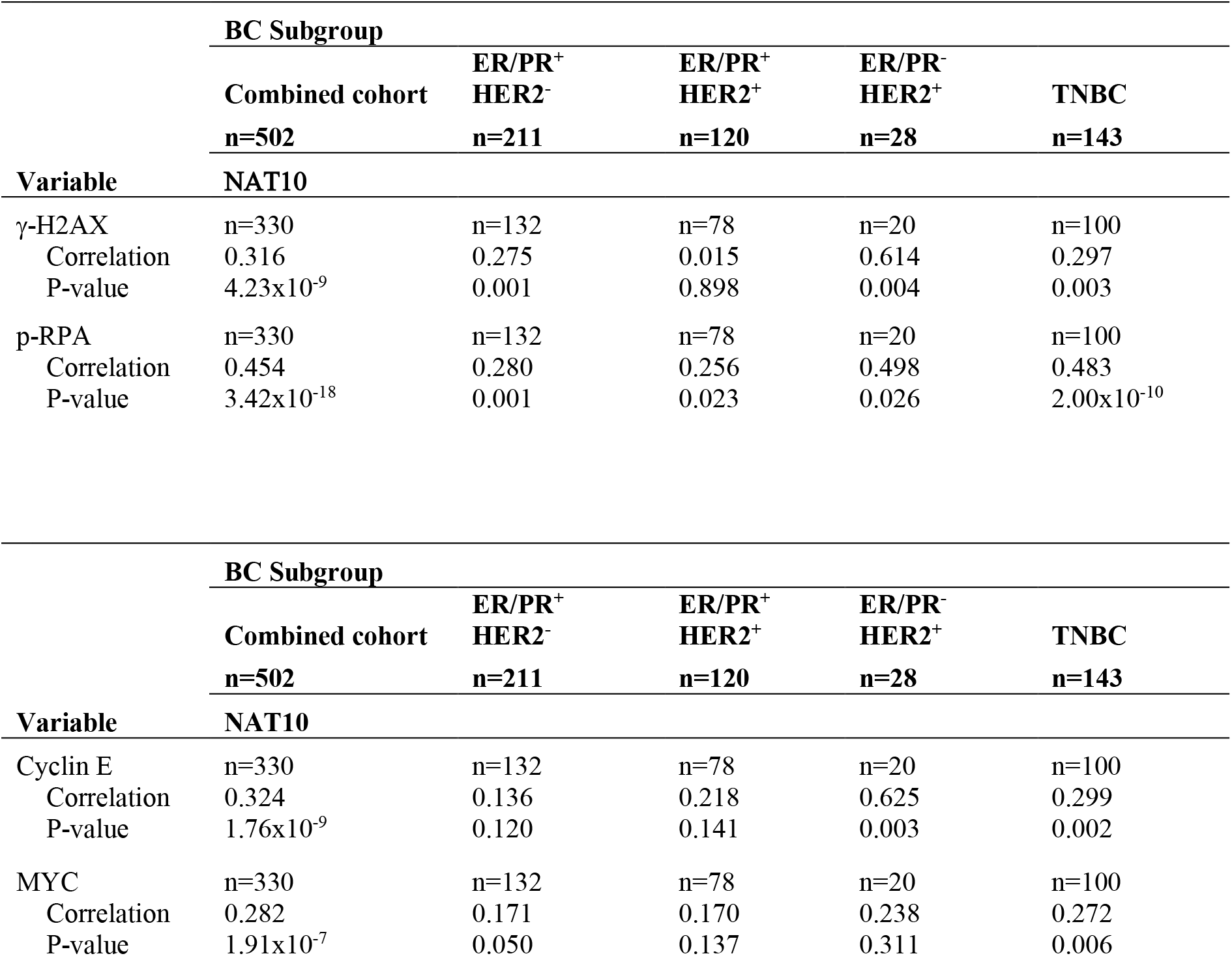
Spearman rank correlation analysis of γH2AX, p-RPA, and oncogene expression versus NAT10 expression in the study population. **A** Spearman correlation analysis of γ-H2AX and p-RPA versus NAT10 expression **B.** Spearman correlation analysis of Cyclin E, MYC versus NAT10 expression (**A**) Association analysis between γH2AX, p-RPA and NAT10 expression in the combined cohort (n=330) and in breast cancer subgroups. (**B**) Association analysis between oncogene expression (Cyclin E and MYC) and NAT10 expression in the combined cohort (n=330) and in breast cancer subgroups.

Next, we tested whether NAT10 expression was also correlated to expression of two oncogenes that are frequently amplified in breast cancer, Cyclin E1 and MYC. NAT10 expression was positively correlated to expression of both Cyclin E1 (R=0.303, p=1.86×10^−9^) and MYC (R=0.264, p=1.45×10^−7^)(Table 1B). Again, associations were most significant in ER/PR^−^/HER2^+^ and TNBC samples (Table 1B). Combined, these analyses validated NAT10 as part of our oncogene-induced replication stress signature, which is associated with markers of replication stress as well as expression of oncogenes, which are known to induce replication stress.

### Landscape of replication stress across tumor types

To investigate the landscape of oncogene-induced replication stress across cancer types, we used the six-gene signature of oncogene-induced replication stress, as a proxy for oncogene-induced replication stress levels. We applied our signature to RNAseq expression data of 8,862 samples retrieved from TCGA. This dataset represents 27 cancer subtypes as well as non-cancer tissues, and displayed large differences in the oncogene-induced replication stress signature score, with diffuse large B cell lymphoma (DLBCL), ovarian cancer and colorectal carcinoma showing highest scores (Fig. 4A). in line with expectations, normal tissues were among the tissue types with lowest scores (Fig. 4A). These observations are in line with previous reports on these cancer subtypes^16,37,38^. To confirm the landscape of oncogene-induced replication stress levels in an independent dataset, we retrieved microarray mRNA expression data of 13,912 patient-derived samples from the GEO database (Fig. 4B). A high concordance between the tumor type replication stress levels in TCGA and GEO was observed (Pearson R=0.77), indicating that our oncogene-induced replication stress signature captures the level of oncogene-induced replication stress also in a platform-independent fashion.

## DISCUSSION

In the present study, we examined transcriptional changes that go along with oncogene-induced replication stress in cell line models and tumor samples to build a gene expression signature of oncogene-induced replication stress. Analysis of a panel of TNBC and non-transformed RPE1 cell lines, combined with analysis of a large dataset of patient-derived cancer samples, yielded a six-gene signature of oncogene-induced replication stress.

Our mRNA-based signature points towards ovarian cancers, colorectal cancers, DLBCLs, TNBCs, cholangiocarcinomas, and esophageal carcinomas having high levels of replication stress. Among these are cancer subtypes that were previously described as prototypical cancers with high levels of replication stress. Specifically, HGSOCs almost invariably have *TP53* mutations (96%), frequently contain high levels of somatic copy number alterations and structural variations, and often have amplification of *MYC* (>30%) and *CCNE1* (>20%)^39^, both of which have been linked to replication stress and genomic instability^13,40^. Likewise, TNBCs show biological similarities with HGSOC and also frequently contain somatic *TP53* mutations as well as amplification of *MYC* and *CCNE1*^41,42^. In good agreement with these data, our immunohistochemical analysis of TNBC samples revealed high levels of p-RPA and γH2AX, markers of single-stranded DNA and DNA breaks, which have associated with replication stress^36^.

Cholangiocarcinoma was among the highest-ranked cancer subtypes based on our replication stress signature. These bile duct cancers have not previously been linked to genomic features associated with replication stress. However, recent studies described recurrent alterations in the proto-oncogene *CCND1*, cell cycle regulatory gene *CDKN2A* as well as the chromatin remodelers *ARID1A*, *IDH1/2* and *PBRM1*^43^, which could underlie DNA replication perturbations^26,44–47^. Of note, mixed type hepatocellular-cholangiocarcinoma was described to share genetic features with hepatocellular carcinoma^48^, including *CCNE1* amplification, which are causally implicated in hepatocellular carcinogenesis^14^.

Also, diffuse large B-cell lymphoma (DLBCL) showed a high score using our gene expression signature. This finding is in line with observations of *MYC* amplification, *ARID1A* mutation and *CDKN2A/B* deletion in DLBCL, with accompanying sensitivity to inhibitors of replication checkpoint kinases^16,49,50^. Although colorectal cancer (CRC) is not commonly regarded as a tumor type with high levels of oncogene-induced replication stress, CRC scored high in our classifier. This observation is strengthened by earlier observations that CRC subgroups in which pRB is inactivated show activation of the DNA damage response^11,51^. Additionally, the chromosomal instability that characterizes CRCs was shown to be linked to replication stress, as judged by slower replication fork progression, and increased numbers of ultrafine anaphase bridges and 53BP1 bodies in G1 cells^37^.

A gene expression signature may aid in patient selection for drugs that target cancers with high levels of replication stress. Early clinical trials evaluating inhibitors of the ATR, WEE1, and CHK1 checkpoint kinases have shown promising results^18,52,53^, but not all patients respond and biomarkers defining optimal patient subgroups has been challenging. For instance, *TP53* mutation status was used to select patients for Wee1 inhibitor treatment, but additional features are needed to define patients who will likely benefit^18,54^. Interestingly, *CCNE1* amplification was among the genetic features that appeared enriched in patients responding to Wee1 inhibitor treatment^18^. Focused analysis of oncogene-induced replication stress in these clinical trials is warranted to test whether a more optimal patient selection is achievable.

## Materials and methods

### Cell lines

hTERT-immortalized human retina epithelial (RPE1), Human embryonic kidney 293 (HEK-293T) and triple-negative breast cancer cell lines MDA-MB-231, BT549, and HCC1806 were obtained from ATCC (#CRL4000, #CRL3216, #HTB26, #HTB122, #CRL2335). RPE1, HEK-293T, and MDA-MB-231 cells were maintained in Dulbecco’s Minimum Essential Media (DMEM, Thermofisher), supplemented with 10% (v/v) fetal calf serum (FCS) and 1% penicillin/streptomycin (Gibco). BT-549 and HCC1806 were cultured in Roswell Park Memorial Institute medium (RPMI, Thermofisher), complemented with 10% FCS and 1% penicillin/streptomycin. All cells were grown at 37°C in normoxic conditions (20% O^2^ and 5% CO^2^). To inactivate *TP53* in RPE1 cells, exon 4 of *TP53* was targeted with CRISPR-CAS9 using the following single guide RNA (sgRNA) (5’-CTGTCATCTTCTGTCCCTTC-3’) and *TP53* mutation was confirmed using Sanger sequencing. Additional technical details are provided in *Supplemental Methods*.

### DNA fiber analysis

To assess replication dynamics, cell lines were pulse-labeled with CldU and IdU for 20 minutes each. Cells were lysed on top of a microscope slides and DNA fibers were spread by tilting the microscope slide and CldU was detected with rat anti-BrdU (Abcam, ab6326), whereas IdU was detected with mouse anti-BrdU (BD Biosciences, Clone B44). The lengths of 300 IdU tracts were measured per condition using ImageJ software. Statistical analysis was performed using the nonparametric Mann-Whitney U test. Additional technical details are provided in *Supplemental Methods*.

### RNA sequencing

Expression of CDC25A, CCNE1, or MYC was induced using 1 μg/mL of doxycycline in biological replicates. Cells were harvested and frozen at −80°C at 48 hours and 120 hours after doxycycline induction. RNA was isolated using the mirVANA kit (Ambion, AM1561). The QuantSeq RNAseq 3’ mRNA kit (Lexogen) was employed to generate cDNA libraries for next-generation sequencing (NGS). The libraries were sequenced with 65 base-pair reads on a NextSeq 500 sequencer (Illumina). Additional technical details are provided in *Supplemental Methods*.

### Immunohistochemistry

Immunohistochemical analysis was performed on tissue specimens taken before treatment of 558 patients with breast cancer who underwent surgery at the University Medical Center Groningen (UMCG) as described previously^36^. Immunohistochemistry was performed with primary antibody against NAT10 (1:500; rabbit, #ab194297, clone EPR18663; Abcam, Cambridge, UK) for 1 hour at room temperature. NAT10 expression was categorized according to percentages of cells that showed staining and on the intensity of staining. Staining intensity was scored in three categories: 0 (negative), 1 (medium), and 2 (high). To calculate the score for each core, the percentage of cells in each group was multiplied by their intensity score, resulting in a range from 0 to 200 points. In addition to NAT10 expression scores, the tumors were scored for the presence of nucleolar NAT10 localization. Next, the scores from each case and staining were averaged and considered for analysis. The status of ER, PR, and HER2 was determined according to the guidelines of the American Society of Clinical Oncology/College of American Pathologists by counting at least 100 cells. Of 558 cases, 410 cases contained complete clinical data and evaluable immunohistochemical staining for NAT10 expression and were included for statistical analysis. Analyses were performed on the 410 cases as well as on patient subgroups based on hormone receptor status and HER2 expression. Additional technical details are provided in *Supplemental Methods*.

## Data availability

All sequencing data have been deposited at the GEO archive of NCBI.

## Acknowledgments

We thank members of the Medical Oncology laboratory for fruitful discussions.

