## Supplemental data for "An mRNA expression-based signature for oncogene-induced replication-stress"

-Supplementary methods
-Supplementary figure legends
-Supplementary figures

### Supplemental methods

#### *Mutagenesis*

*TP53* was mutated in RPE1 cells as described previously[1]. In short, exon 4 of *TP53* was targeted with CRISPR-CAS9 using the following single guide RNA (sgRNA) (5'-CTGTCATCTTCTGTCCCTTC-3'), cloned into pSpCas9(BB)-2A-GFP, which was a gift from Feng Zhang[2] (PX458, plasmid #48138, Addgene). Subsequently, PX458 was transfected into RPE1 cells, followed by three weeks of selection with Nutlin-3a (10 µM, Axon Medchem). The resulting RPE1-*TP53*$^{mut}$ cells were sorted into a monoclonal cell line using a MoFLO XDP cell sorter. *TP53* mutations in exon 4 were confirmed by Sanger sequencing, and lack of p53 expression was confirmed by Western blot analysis. One *TP53* allele was mutated in exon 4 by a 7 base pair deletion (TCA-TCT-T), leading to a frameshift at codon 97. The other *TP53* allele showed a 217 base pair insertion, also leading to a frameshift.

#### *DNA cloning and Retroviral infections*

RPE1-*TP53*$^{wt}$, RPE1-*TP53*$^{mut}$, MDA-MB-231, BT549, and HCC1806 cell lines were stably transduced with pRetroX-Tet-On Advanced (Clontech). To this end, HEK-293T cells were transfected with ten µg of pRetroX-Tet-On Advanced, 2.5 µg of pMDg, and 7.5 µg of pMD-g/p as described previously[3]. At 24, 36, and 48 hours after transfection, virus-containing supernatant was filtered and added to target cells, which were subsequently selected for seven days using geneticin (G418 Sulfate, 800 µg/mL, Thermofisher). Next, target cell lines harboring pRetroX-Tet-On Advanced were transduced with pRetroX-Tight-Pur containing *CDC25A*, *CCNE1* or *MYC*. To this end, human *CDC25A* was PCR amplified from FLAG-CDC25A-WT, which was a gift from Peter Stambrook[4], using the following oligos: forward: 5'-CGCGGCCGCCATGGAACTGGGCCCGGAGCCC-3', reverse: 5'-GATGAATTCTCACAGCTTCTTCAGACG-3'. Human *CCNE1* was PCR amplified from Rc-CycE, which was a gift from Bob Weinberg (Plasmid #8963, Addgene)[5], using the following oligos: forward: 5'-CGCGGCCGCCATGAAGGAGGACGGCGGCGCG-3', reverse: 5'-GATGAATTCTCACGCCATTTCCGGCCC-3'. The resulting fragments were cloned into pJET1.2/blunt, GeneJET, (ThermoFisher). Human *MYC* was PCR amplified from MSCV-MYC -T58A-puro, which was a gift from Scott Lowe (Plasmid #18773, Addgene)[6], using the following oligos: forward: 5'-CGCGGCCGCCATGCCCCTCAACGTTAGCTTC-3', reverse: 5'-GATGAATTCTTACGCACAAGAGTTCCG-3'. *CDC25A*, *CCNE1*, and *MYC* were subcloned into pRetroX-Tight-Pur using NotI and EcoRI restriction sites. Subsequently, cell lines harboring pRetroX-Tet-On Advanced were transduced with pRetroX-Tight-Pur containing *CDC25A*, *CCNE1*, *MYC*, or empty plasmid. After three rounds of transduction, cells were selected for two days with 1 µg/mL (MDA-MB231, BT-549, and HCC1806) or 5 µg/mL (RPE1) of puromycin dihydrochloride (Sigma).

***Western blotting***

Cells were lysed in M-PER lysis buffer (Pierce), supplemented with protease and phosphatase inhibitor cocktail (Thermo Scientific). Protein content was measured using the Pierce BCA protein quantification Kit (Thermo Scientific). Protein samples were separated using SDS-polyacrylamide gels (SDS-PAGE) and transferred to polyvinylidene fluoride membranes (Immobilon). Membranes were blocked in 5% skimmed milk (Sigma) in Tris-buffered saline (TBS) containing 0.05% Tween-20 (Sigma) and incubated overnight with primary antibodies at 4ºC and subsequently incubated with secondary antibodies for 1 hour at room temperature. Primary antibodies used were mouse anti-MYC (Santa Cruz Biotechnology, Sc-40, 1:200), mouse anti-CDC25A (Santa Cruz Biotechnology, Sc-7389, 1:200), mouse anti-CCNE1 (Abcam, ab3927, 1:500), mouse anti-p53 (Santa Cruz Biotechnology, Sc-126, 1:1,000) and mouse anti-β-actin (MpBiomedicals, 69100, 1:10,000). Secondary antibodies used were horseradish peroxidase-linked anti-mouse IgG (1:2,000, DAKO) and visualized using chemiluminescence (Lumi-Light, Roche Diagnostics) on a Bio-Rad bioluminescence device. Protein imaging was performed using Image Lab software (Bio-Rad).

***Immunofluorescence microscopy***

RPE1-$TP53^{wt}$ or RPE1-$TP53^{mut}$ doxycycline-inducible cell lines were seeded on glass coverslips. Cells were treated with doxycycline (1 µg/mL) for 48 hours or left untreated. Next, cells were fixed in 3.7% formaldehyde in PBS for 15 minutes and permeabilized in 0.1% Triton X-100 for 5 minutes. Subsequently, cells were incubated with primary and secondary antibodies at room temperature for 80 minutes, respectively. Primary antibodies used were mouse anti-MYC (Santa Cruz Biotechnology, Sc-40, 1:100), mouse anti-CDC25A (Santa Cruz Biotechnology, Sc-7389, 1:50 or 1:100) and mouse anti-CCNE1 (Abcam, ab3927, 1:100). Secondary antibody used was Alexa488-conjugated rabbit anti-mouse (1:300), and DAPI was applied as counterstaining. Images were acquired on a Leica DM-6000B (63x immersion objective with 1.30 NA) or DM-4000B (63x immersion objective with 0.5 NA) fluorescence microscope, equipped with Leica Application Suite software.

#### *DNA fiber analysis*

To assess replication dynamics, RPE1-*TP53*$^{wt}$, RPE1-*TP53*$^{mut}$, MDA-MB-231, BT-549, and HCC1806 doxycycline-inducible cell lines were pulse-labeled with CldU (25 µM) for 20 minutes. Next, cells were washed with warm medium and pulse-labeled with IdU (250 µM) for 20 minutes either alone or in combination of hydroxyurea (0.1 or 5 mM). Cells were collected using trypsin and lysed on top of a microscope slide in lysis solution (0.5% sodium dodecyl sulfate (SDS), 200 mM Tris [pH 7.4], 50 mM EDTA). DNA fibers were spread by tilting the microscope slide and were subsequently air-dried and fixed in methanol/acetic acid (3:1) for 10 minutes. Next, DNA spreads were immersed in 2.5M HCl for 75 minutes. DNA fibers were blocked in blocking solution (5% BSA in PBS) for 30 minutes and incubated with primary antibodies for 60 minutes at room temperature. CldU was detected with rat anti-BrdU (1:1,000, Abcam, ab6326), whereas IdU was detected with mouse anti-BrdU (1:250, BD Biosciences, Clone B44). Secondary antibodies used for detection were Alexa488-conjugated goat anti-rat IgG (1:500) and Alexa647-conjugated goat anti-mouse IgG (1:500). Images were acquired on a Leica DM-6000B (63x immersion objective with 1.30 NA) fluorescence microscope equipped with Leica Application Suite software. The lengths of 300 IdU tracts were measured per condition using ImageJ software. Statistical analysis was performed using the nonparametric Mann-Whitney U test in GraphPad Prism 6.

#### *RNA sequencing*

Expression of CDC25A, CCNE1, or MYC in MDA-MB-231, BT549, HCC1806, RPE1-*TP53*$^{wt}$, and RPE1-*TP53*$^{mut}$ cells was induced using 1 µg/mL of doxycycline. Cells were harvested and frozen at -80°C at 48 hours and 120 hours after doxycycline induction. Next, RNA was isolated using the mirVANA kit (Ambion, AM1561). To determine RNA quality, RNA was separated by electrophoresis on microfluidic sipper chips and detected by fluorescence (LabChip GX, Caliper LifeSciences). RNA Quality Scores (RQS) were based on electropherogram features (total and fast region areas, 28S, and 18S height), and ranged from 0-10. Only samples with RQS scores above 5 were included for analysis. To generate cDNA libraries suitable for next-generation sequencing (NGS), the QuantSeq RNAseq 3' mRNA kit (Lexogen) was employed. Briefly, the poly-A tail from total RNA was bound to oligo-dT primers to synthesize the first cDNA strand. After RNA removal, a second cDNA strand was synthesized by random priming. The double-stranded cDNA library was purified, and PCR amplified with Illumina sequencing adapters. The libraries were sequenced with 65 base-pair reads on a NextSeq 500 sequencer (Illumina) and generated 7.2 to 19.8 million of reads per sample. FastQC and Samtools Flagstat software assessed RNA sequencing quality control according to manufacture guidelines.

***Quality control and normalization***

*Cell line RNA-sequencing data*

Samples from doxycycline-treated cells expression oncogenes at 48 and 120 hours were analysed as a pooled group of samples with oncogene-induced replication stress. Ensembl IDs having a robust average read count (Hodges Lehmann estimate) lower than 20, were removed from the analysis. Thereafter, normalization was performed on the RNA sequencing data. Firstly, the difference in read counts due to difference in cell lines was adjusted for every cell line separately (MDA-MB-231, BT549, HCC1806, RPE-1-*TP53*$^{wt}$ and RPE-1-*TP53*$^{mut}$) using the following steps:

1. The robust average read count for each Ensembl ID was obtained using Hodges Lehmann estimator.
2. The robust standard deviation of read counts for each Ensembl ID was obtained using Hall's estimator.
3. The read count of each Ensembl ID was normalized using the following formula

*Adjusted read count = (read count – robust average)/robust standard deviation*

To eliminate difference between read counts of doxycycline-treated samples and non-doxycycline-treated samples, the following steps were taken:

4. The robust average read count corresponding to doxycycline-treated samples in the empty plasmid for each Ensembl ID was obtained using Hodges Lehmann estimator (RA_empty_dox).
5. The robust average read count corresponding to non-doxycycline induced samples in the empty plasmid samples for each Ensembl ID was obtained using Hodges Lehmann estimator (RA_empty_no_dox).
6. The robust standard deviation of read counts corresponding to doxycycline induced samples in the empty plasmid for each Ensembl ID was obtained using Hall's estimator (RSD_empty_dox).
7. The robust standard deviation of read counts corresponding to non-doxycycline treated samples in the empty plasmid for each Ensembl ID was obtained using Hall's estimator (RSD_empty_no_dox).
8. The read counts for all doxycycline-treated samples were adjusted using the following formula:

*Adjusted read count = (read count - RA_empty_dox)/ RSD_empty_dox*

9. The read counts for all non-doxycycline induced samples were adjusted using the following formula:

   *Adjusted read count = (read count - RA_empty_no_dox)/ RSD_empty_no_dox*

***TCGA RNA-sequencing data***

Genes having robust average read counts (Hodges Lehmann estimate) lower than 20, were removed from the analysis. Thereafter, normalization was performed on the RNA sequencing data. To this end, the difference in read counts due to difference in cancer types was adjusted for every cancer type separately using the following steps:

1. The robust average read count for each gene was obtained using Hodges Lehmann estimator.
2. The robust standard deviation of read counts for each gene was obtained using Hall's estimator.
3. The read count of each gene was normalized using the following formula

   *Adjusted read count = (read count – robust average)/robust standard deviation*

***Permutational multivariate analysis of variance (PERMANOVA)***

PERMANOVA was conducted on the mRNA expression dataset after adjusting for cell line-specific effects and treatment with doxycycline to analyse differences between means of gene expression values for overexpressed/amplified samples compared to normal/neutral samples for the corresponding oncogene. This method aims to perform classical partitioning with no assumption of multivariate normality. The model for gene *g* corresponding to samples with over-expression of oncogene *o* is the following:

$RNAseq\_expression_{g,o} = alpha_{g,o} + beta_{g,o} * oncogene\_induction\_indicator_{o}$

In this model, $oncogene\_induction\_indicator_{o}$ = 1, if in the sample the oncogene *o* is induced, otherwise this value = 0

***Classification of TCGA samples based on copy number data***

For each oncogene *o*, all TCGA samples are categorized into three groups ('amplified', 'neutral' or 'deleted'). Samples with a log2(copy number/2) value of each oncogene *o* smaller than -0.2 were categorized as 'deleted' ($D_o$). Samples with a copy number variation value of each oncogene greater than 0.2 were categorized as 'amplified' ($A_o$). Samples with a copy number variation value of each oncogene between -0.2 and 0.2 were categorized as ‘neutral’ ($N_o$). Differential mRNA expression analysis based on oncogene classification was done

For every oncogene *o*, differential mRNA expression analysis was conducted between samples of class $A_o$ and $N_o$ using the following steps:

1. For every gene *g*, which followed quality control criteria
2. Mean of mRNA expression values of gene g for samples in class $A_o$ were compared to the mean of mRNA expression values of gene g for samples in class $N_o$ using Welch’s t-test with unequal variance.
3. A metric ($metric_{g,o}$) was obtained from the Welch’s t-test as $-log_{10}$(P-value)*sign(t statistic).

***Sensitivity of differential mRNA expression analysis***

Classification of samples into three groups (deleted, neutral and amplified) were conducted on multiple cut-off values of log2(copy number/2) to check the sensitivity of differential mRNA expression analysis. Cut-off values used for categorizing deleted samples were -0.2, -0.21, -0.22, -0.23, -0.24, -0.25, -0.26, -0.27, -0.28, -0.29, -0.3. Cut-off values used for categorizing amplified samples were 0.2, 0.21, 0.22, 0.23, 0.24, 0.25, 0.26, 0.27, 0.28, 0.29, 0.3. Correspondingly neutral samples were also categorized using the above cut-off values. Along with these cut-off values, per sample different cut-off values were obtained using GISTIC2.0 algorithm.

***Gene Set Enrichment Analysis***

GSEA was performed utilizing 12 gene set databases from the MSigDB.17 Gene sets containing less than 10 genes or more than 500 genes (after filtering out genes that were not present in our data sets) were excluded from further analysis. Enrichment of a gene set was tested according to the two-sample Welch’s t-test for unequal variance.

Welch’s t-test was conducted between the set of metrics from PERMANOVA or Welch’s t-test of genes whose corresponding gene identifiers are members of the gene set under investigation and metrics of genes whose corresponding gene identifiers are not members of the gene set under investigation. To be able to compare gene sets of different sizes, Welch’s t statistics were transformed to z-scores. To control the false discovery rate, we performed a multivariate permutation test with 100 permutations. For each permutation round gene identifiers were randomly assigned to metrics. This allowed us to present the number of significantly enriched gene sets per gene set database using a false discovery rate of 1% and a confidence level of 80%.

#### *Prediction of gene functionalities*

We used a GBA approach to predict likely functions for genes based on gene co-regulation. For this, we conducted a consensus-ICA on an unprecedented scale (manuscript under review). In short, a covariance matrix was calculated between 19,635 genes using the expression patterns of 106,462 gene expression profiles generated with Affymetrix HG-U133 Plus 2.0 representing the many disease states, cellular states, and genetic and chemical perturbations that were obtained. Consensus-ICA was performed on the covariance matrix, which resulted in the identification of a large set of CESs and a mixing matrix reflecting the activity of each source in the expression pattern of each gene across the samples. Next, a GBA approach was used to predict the functionality of individual genes. First, we retrieved 16 public gene set collections describing a large range of biological processes and phenotypes. For each gene set, we calculated its 'bar code' by averaging the MM weight of its member genes. Next, for each gene in the MM, the distance correlation was determined between its MM weights and the gene set bar code. A high correlation between a gene's MM weight and a gene set bar code indicated that the gene under investigation shared a functionality with the genes of the specific gene set under investigation. Significance levels were obtained with permutated data (250 permutations). This strategy was used on 23,372 well-described functional gene sets, which enabled us to create a comprehensive network of predicted functionalities of individual genes. This framework is available at http://www.genetica-network.com.

#### *Immunohistochemistry*

Immunohistochemical analysis was performed on tissue specimens taken before treatment of 558 patients with breast cancer who underwent surgery at the University Medical Center Groningen (UMCG) as described previously[7].

TMA slices were deparaffinized using xylene. Antigen retrieval was done using microwave treatment in Tris/EDTA buffer (pH 9) for 15 minutes. Slides were incubated for 30 minutes in 0.3% hydrogen peroxidase ($H_2O_2$) to suppress endogenous peroxidase activity. Immunohistochemistry was performed with primary antibody against NAT10 (1:500; rabbit, #ab194297, clone EPR18663; Abcam, Cambridge, UK) for 1 hour at room temperature. Subsequently, slides were incubated with horse-radish-peroxidase (HRP)-conjugated secondary antibody goat anti-rabbit (1:100, Dako) for 30 minutes. Staining was detected by the application of 3,3-diaminobenzidine (DAB), and hematoxylin as a counterstaining. Scoring was performed semi-quantitatively by a researcher and was supervised by a breast cancer pathologist. NAT10 expression was categorized according to percentages of cells that showed staining and on the intensity of staining. Staining intensity was scored in three categories: 0 (negative), 1 (medium), and 2 (high). To calculate the score for each core, the percentage of cells in each group was multiplied by their intensity score, resulting in a range from 0 to 200 points. In addition to NAT10 expression scores, the tumors were scored for the presence of nucleolar NAT10 localization. Next, the scores from each case and staining were averaged and considered for analysis. The status of ER, PR, and HER2 was determined according to the guidelines of the American Society of Clinical Oncology/College of American Pathologists by counting at least 100 cells.

Immunohistochemical stainings were considered evaluable when a tumor core contained at least 10% of tumor cells. Also, tumor stainings were included for analysis when at least 2 out of 3 cores were evaluable. Core loss over 558 cases was 20.4% (NAT10 expression, n=114; NAT10 nucleoli, n=114). Of 558 cases, 410 cases contained complete clinical data and evaluable immunohistochemical staining for NAT10 expression and NAT10 nucleoli and were included for statistical analysis. Analyses were performed on the 410 cases as well as on patient subgroups based on hormone receptor status and HER2 expression. Numbers of patients in in each subgroup were as follows: ER/PR$^+$HER2$^-$ (n=164), ER/PR$^+$HER2$^+$ (n=95), ER/PR$^-$HER2$^+$ (n=21) and TNBC (n=130). Differences regarding immunohistochemical expression levels or nucleoli amounts between the four groups were analyzed using Mann-Whitney U tests. Associations between different immunohistochemical expression levels or nucleoli amounts were calculated using Spearman correlation tests. All statistical analyses in this study were performed using SPSS Statistics 23.0 (IBM).

**Supplementary figure legends**

**Supplementary Figure 1:** ***Overexpression of CDC25A, CCNE1, or MYC upon doxycycline treatment.*** **(A-D)** Immunoblotting of CDC25A, CCNE1, MYC, p53, and β-Actin after 48 hours after doxycycline treatment of indicated RPE1-*TP53*$^{mut}$, MDA-MB-231, BT549, and HCC1806 cell lines.

**Supplementary Figure 2:** ***Immunohistochemical NAT10 analysis breast tissue.***
**(A)** Representative staining of NAT10 and IgG control staining in breast tissue. Scale bar in the left panel is 1 mm, whereas in the right panel represents 10 μm.

### Suppl. Figure 1

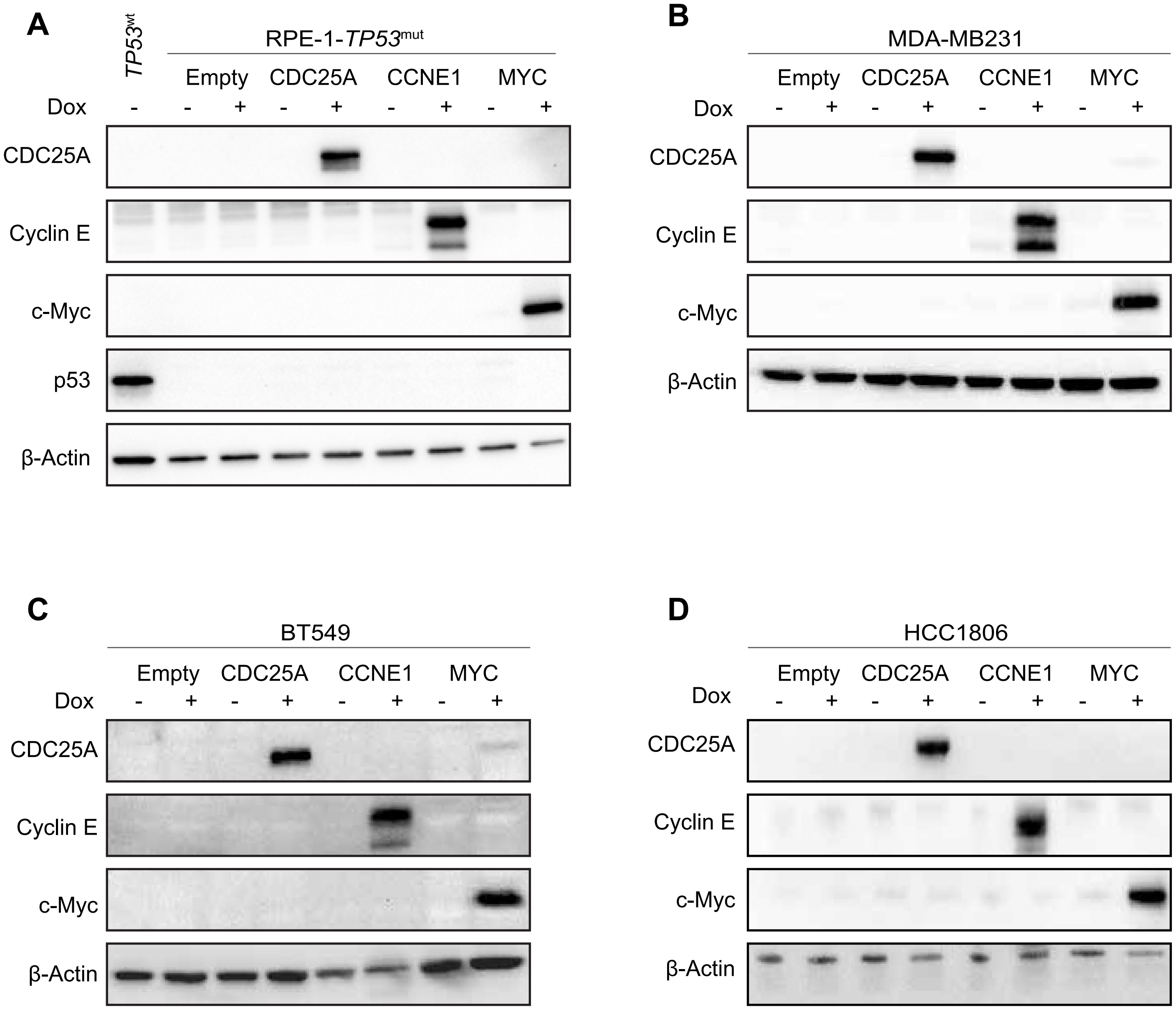

### Suppl. Figure 2

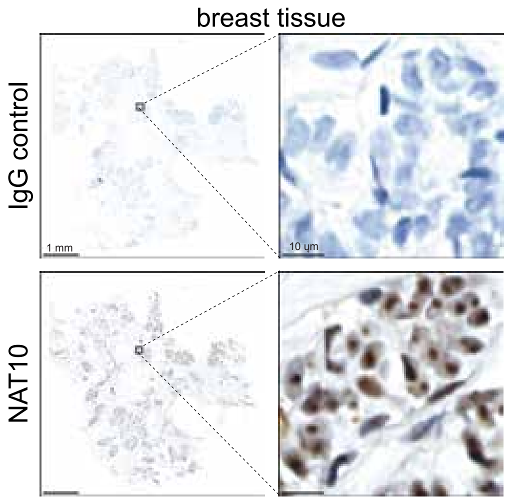
